# Longevity or Well-being? A Dual-Dimension Structure of Neuroticism

**DOI:** 10.1101/2024.07.23.604876

**Authors:** Yini He, Jing Xiao, Ke Hu, Tian Gao, Yan Yan, Lei Wang, Kaixin Li, Wenkun Lei, Kun Zhao, Changsheng Dong, Xiaohan Tian, Chaoyue Ding, Yingjie Peng, Junxing Xian, Shangzheng Huang, Xiya Liu, Long Li, Peng Zhang, Zhanjun Zhang, Sheng He, Ang Li, Bing Liu

## Abstract

The development of personality traits is often viewed as evolutionarily adaptive. Current neuroticism research, however, predominantly highlights its negative health impacts, neglecting its potential evolutionary advantages. We propose that neuroticism’s inter-individual variability can be structured into two distinct geometric dimensions. One, named the Emotional Reactivity-Instability/Distress Spectrum (ERIS), correlates strongly with longevity and is associated with chronic diseases and risk-averse lifestyle. This dimension is underpinned by evolutionarily conserved subcortical brain regions and genes. The other, resembling the overall neuroticism score, is primarily linked to mental and stress-related disorders, as well as life satisfaction. It involves higher-order emotional brain regions and is genetically enriched in human-accelerated regions. Collectively, these dimensions represent a dual-strategy personality framework that optimizes survival and well-being, with the former being evolutionarily conservative and the latter potentially a unique human adaptation.

## Introduction

Extensive empirical evidence has established neuroticism as a fundamental domain of human personality^1,2^. Neuroticism describes an individual’s general tendency towards experiencing negative emotions, such as worry, depression, irritability, feelings of helplessness, and mood instability^3,4^.

Given that neurotic individuals have poor responses to environmental stress, studies over the past decades have highlighted its negative impact on public health^5,6^. High levels of neuroticism are associated with various negative outcomes, such as susceptibility to mental and physical disorders^7,8^, diminished quality of life^5,6,8^, and increased mortality risk^9,10^. However, from an evolutionary standpoint, minimal reactions to threatening stimuli, akin to an extreme form of low neuroticism, are generally not advantageous for survival. This concept echoes the ancient Chinese adage, ‘*Life springs from sorrow and calamity; death comes from ease and pleasure*’, encapsulating a time-honored survival philosophy. To mitigate risks and ensure survival, both animals and human ancestors required automatic responses to immediate and potential future threats^11,12^. This necessity is manifested through adaptive emotions such as fear and its anticipatory form -- anxiety, which are supported by distributed brain regions, such as the hippocampus, amygdala, and medial prefrontal cortex^12–14^.

Neuroticism presents a significant paradox in the field of personality psychology: epidemiologically, it is associated with detrimental health outcomes, including increased mortality rates, yet it may simultaneously confer evolutionary advantages in survival scenarios. While certain studies have acknowledged heterogeneity within neuroticism^9,15–17^, the complexity of this paradox remains largely unresolved. This obscurity impedes our understanding of the nature and origins of neuroticism and hampers the development of effective intervention strategies^2^. **In this study, we propose that neuroticism has evolved distinct dimensions, potentially as adaptations to unique ecological and cultural changes, influencing lifestyle and health outcomes through diverse genetic and neural mechanisms.** Beyond the psychometric approach, we developed an inter-subject network to comprehensively characterize individual differences in neuroticism and utilized a dimensionality reduction technique to delineate the major gradients of population structure. By analyzing neuroticism questionnaires, electronic health records, behavioral phenotypes, neuroimaging, and genetic information in the UK Biobank, we reveal the heterogeneous gradients of neuroticism across populations and demonstrate how these gradients differently correlate with health outcomes, disease susceptibility, and lifestyle factors, as well as neurobiological and genetic underpinnings. We posit that an enhanced understanding of these gradients of neuroticism could be invaluable in formulating more effective preventive health strategies and advancing public health initiatives.

## Results

### Robust dual-dimensional structure of neuroticism across diverse populations

To systematically examine the latent structure of neuroticism, we analyzed five independent datasets across different cohorts. Four of the datasets were based on the neuroticism subscale of the NEO Five-Factor Inventory (FFI-N)^18^(Fig. 1a, left): the Human Connectome Project^19^ (HCP, *n*=1198, adults, United States), the Human Connectome Project Development^20^ (HCP-D, *n*=229, adolescents and young adults, United States), the Chinese university student dataset (CN-U, *n*=612, adults, China), and the Chinese adolescent dataset (CN-A, *n*=531, adolescents, China). Additionally, one of the datasets utilized the neuroticism subscale of the Eysenck Personality Questionnaire-Revised Short Form (EPQ-N)^21^(Fig. 1a, right), specifically the UK Biobank^22^ (*n*=401,574 elder adults, United Kingdom). For better interpretation and visualization, we categorized each item into ‘emotional reactivity’ and ‘emotional instability/distress’ (see Methods) but refrained from any quantitative analysis based on this classification. As illustrated in Fig. 1b, instead of a focus solely on total scores, we constructed an inter-subject similarity network based on neuroticism item-level scores and employed a diffusion map^23^ approach to derive individual low-dimensional embeddings (see Methods). Recently, such an embedding approach has also been applied to characterize the dominant axes of the inter-subject brain similarity network^24^. Utilizing this technique, we uncovered a topological structure of neuroticism spanning the continuous spectrum, leading to the identification of two interpretable neuroticism gradients across five datasets (see below). As described in Supplementary Note 1, Supplementary Table 1-5, exploratory factor analysis did not consistently yield uniform latent structures across different questionnaires and cohorts.

**Fig. 1.**
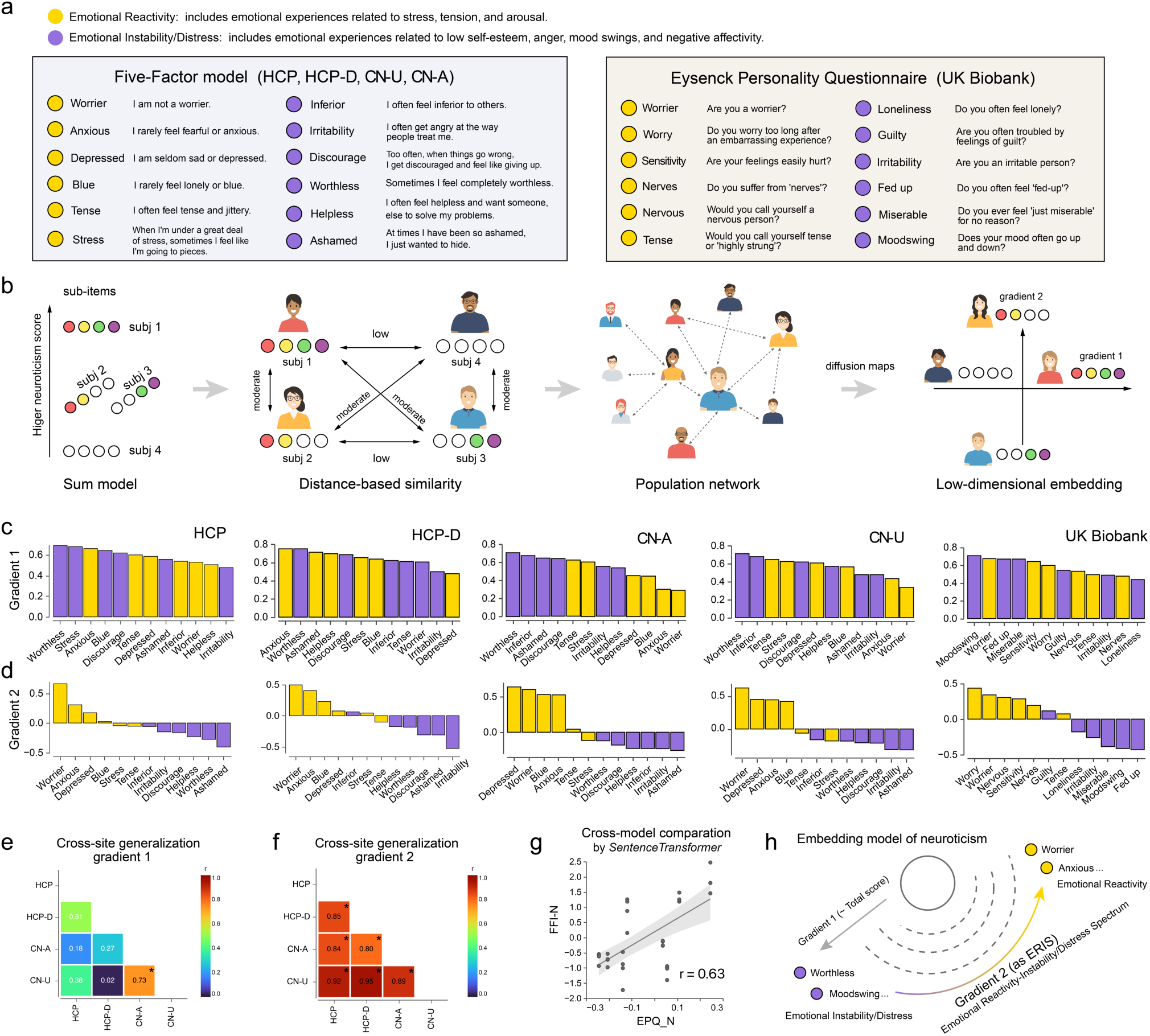
Geometric structure of neuroticism. **(a)** The assessment of the neuroticism trait employs scales from both the Neuroticism subscale of the NEO Five-Factor Inventory (FFI-N) and the Neuroticism subscale of the Eysenck Personality Questionnaire-Revised Short Form (EPQ-N). Detailed questions from these subscales are categorized into ‘emotional reactivity’ and ‘emotional instability/distress’ (see Methods). To visually distinguish these categories, they are represented in lavender and ocher colors, respectively. **(b)** The schematic illustrates our pipeline for deriving geometric embeddings/gradients of neuroticism. This is achieved by constructing a between-subject similarity network based on the similarity of items, followed by employing diffusion maps to obtain individual-level gradient scores. As an example within this framework, it’s possible for individuals to have identical overall neuroticism scores, like subjects 2 and 3, yet possess distinctly different scores on gradient 2. **(c-d)** The weights of items contributing to the principal and second neuroticism gradients are displayed across five varied datasets (from left to right: HCP, HCP-D, CN-U, CN-A, and UK Biobank). The vertical axis quantifies the weight of each item on the gradient 1(c) and gradient 2(d), reflecting the respective contribution of each item to the gradient (Spearman correlation coefficient between the original item and gradient scores across population). Concurrently, the horizontal axis lists the specific item questions as outlined in section a, with the color scheme directly corresponding to the categorizations established therein. **(e-f)** The cross-cohort analysis utilizes FFI-N across datasets to assess the stability of item weights for gradient 1 (e) and gradient 2, marked by an asterisk for unadjusted *P* values below 0.05 (n=12, across the items). **(g)** Sentence transformer models indicate item-level concordance between the FFI-N and the EPQ-N, specifically for gradient 2 of neuroticism. **(h)** Drawing on the style of the earlier brain’s functional gradient schematic^23^, we depicted the geometric structure of neuroticism based on our findings. Importantly, we identified and characterized gradient 2 as emotional reactivity-instability/distress spectrum (ERIS), encapsulating a continuous spectrum dimensional phenotype ranging from emotional reactivity to instability/distress.

The principal gradient statistically approximates the conventional total neuroticism score (HCP: *r* = 0.997, *P* < 2.2×10^-308^; HCP-D: *r* = 0.998, *P* < 2.2×10^-308^; CN-U: *r* = 0.994, *P* < 2.2×10^-308^; CN-A: *r* = 0.990, *P* < 2.2×10^-308^; UK Biobank: *r* = 0.994, *P* < 2.2×10^-308^; age-and sex-adjusted Spearman’s correlations), suggesting that the largest variance across population network is a composite reflection of all neuroticism items (Fig. 1c). Intriguingly, the second gradient unveils a novel geometric dimension of neuroticism. This dimension bifurcates into positive emotional reactivity, exemplified by anxiety, worry, tension, and depressed mood, as well as negative emotional instability/distress, characterized by irritability, shame, and mood instability (Fig. 1d). These two embeddings of neuroticism exhibit no significant correlation across individuals (HCP: *r* = 0.04, *P* = 0.13; HCP-D: *r* = -0.007, *P* = 0.91; CN-U: *r* = -0.05, *P* = 0.23; CN-A: *r* = 0.02, *P* = 0.63; UK Biobank: *r* = -0.003, *P* = 0.08; age-and sex-adjusted Spearman’s correlations). Within the datasets using FFI-N, the second gradient demonstrated remarkable consistency in items of item weightings (Fig. 1f), in contrast to the principal gradient, which showed limited item specificity (Fig. 1e). To quantify and compare item weightings between the two types of questionnaires, we utilized a sentence transformer model to vectorize the item’s sentences^25^, suggesting the existence of a shared second gradient across two different questionaries (Fig. 1g, *r* = 0.63, *P* = 1.22×10^-4^, see Methods).

Our results suggest that the neuroticism personality trait encompasses a multidimensional geometric structure (Fig. 1h), exhibiting stable patterns across diverse populations, age groups, and measurement methodologies. Specifically, we have identified a stable, secondary dimension of neuroticism that encompasses two distinct aspects based on scoring. Higher scores indicate an increase in anxiety, worry, tension, and depressed mood, which we refer to as emotional reactivity. At the same time, they also reflect a decrease in irritability, frustration, and mood instability, which we refer to as emotional instability or distress. For clarity, we have labeled this gradient the emotional reactivity-instability/distress spectrum (abbreviated as ERIS), and the principal gradient as ‘neuroticism’, analyzing the latter using the total neuroticism score in subsequent analyses. A comprehensive discussion is provided in Supplementary Note 2 to elaborate on the relationship between ERIS and previous studies on the structure of neuroticism.

### The ERIS gradient exhibits a notable survival advantage

While abundant evidence suggests that high levels of neuroticism correlate with higher mortality rates^1,2,26^, recent neuroticism phenotype or genetic studies at the facet and item level have identified that in certain circumstances, high neuroticism may indicate lower mortality rates^9,15–17^. However, to date, no study has directly employed the neuroticism scale to identify a subgroup whose lifespan exceeds that of individuals traditionally considered the healthiest with the lowest levels of neuroticism. As illustrated in Fig. 2a, we categorically segregated the UK biobank participants into four exclusive groups: high and low neuroticism, and high and low ERIS levels, respectively. This classification was rigorously applied within each gender and age category, as detailed in the Methods section. In our survival analysis over more than 15 years, it was observed that individuals in the high ERIS group exhibited significantly lower mortality rates compared to those in the lowest neuroticism group (Fig. 2b, *P* = 1.3×10^-6^). We conducted a comparative analysis using specific factors of neuroticism identified in previous studies, which also suggested a lower mortality rate among individuals in our defined high ERIS subgroup (see Supplementary Note 2, Supplementary Table 6, and Supplementary Fig. 1). Notably, the low ERIS group displayed a marked inclination towards the highest mortality rates, ranking at the bottom among the four groups. This pattern of findings is visually consistent across different age and gender categories (Fig. 2c).

**Fig. 2.**
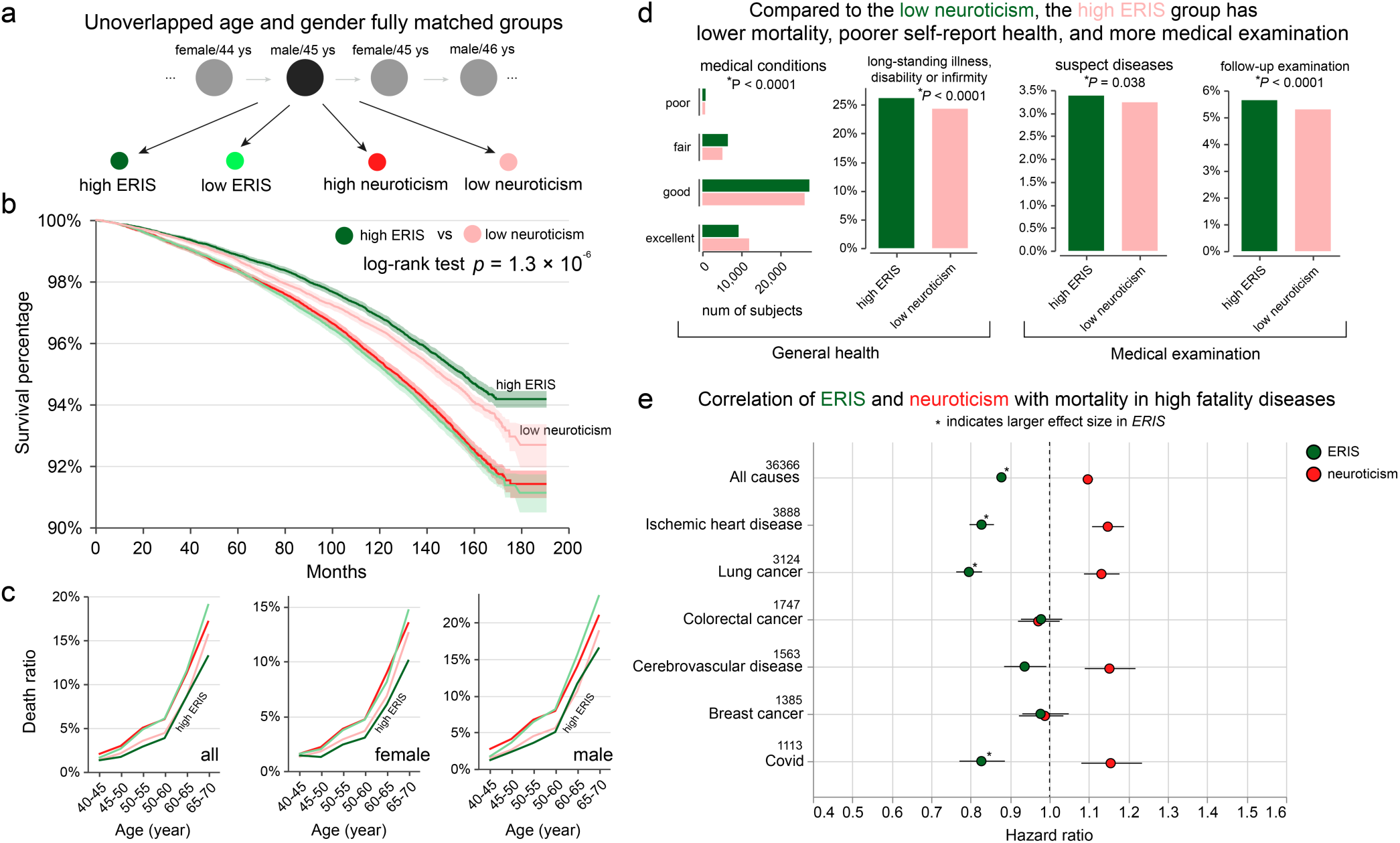
Longitudinal impact of neuroticism gradients on mortality rates. **(a)** Schematic for subgrouping by ERIS and neuroticism, categorizing participants into four distinct groups: low/high neuroticism and low/high ERIS. Duplicates were removed, and group sizes were balanced by adjusting larger groups to match the smallest group, ensuring strict age and gender control. The low neuroticism group comprised solely participants with zero scores on neuroticism. (see Methods). **(b)** Kaplan-Meier survival curves illustrate the survival rates for four extreme groups (grouped according to the method in a), ranked from highest to lowest as follows: high ERIS, low neuroticism, high neuroticism, and low ERIS. For the two groups with higher survival rates, a log-rank test comparison revealed that the high ERIS group’s survival rate was significantly greater than that of the low neuroticism group. **(c)** Stratified death ratios were conducted for extreme ERIS and neuroticism groups, categorized by age and gender. Age intervals were established in five-year increments, with analyses performed both on the total sample and separately for each gender. Across different gender and age comparisons, the high ERIS group generally exhibited the lowest mortality rates. **(d)** Given the lower mortality rates observed in the high ERIS group compared to the low neuroticism group (as illustrated in Fig. b-c), further investigation was conducted into the differences in health self-evaluation and medical care behaviors between individuals with high ERIS and those with low neuroticism. The bar chart indicates that individuals with high ERIS reported poorer self-assessed health and were more likely to undergo medical examinations. The deep green bars represent the high ERIS group, while the light pink bars represent the low neuroticism group. A chi-squared test was used to evaluate the differences in self-reported health and medical examination rates between the two groups, with the p-values indicating statistical significance. General health metrics included self-reported medical conditions and evaluations of long-term illnesses, disabilities, or infirmities; medical examinations encompassed checks for suspected diseases and follow-up examinations after treatments.**(e)** Hazard ratios of Cox proportional hazards regression between ERIS/neuroticism and mortalities including all causes, ischemic heart disease, lung cancer, colorectal cancer, cerebrovascular disease, breast cancer. The corresponding death numbers are presented on top of the disease name. The asterisk denotes that ERIS exhibits a greater effect size than neuroticism, as shown in Supplementary Tab.7.

After establishing lower future mortality rates in the high ERIS group compared to the low neuroticism group, we scrutinized their baseline self-reported health status. Notably, a higher percentage of individuals in the low neuroticism group self-assessed as being in excellent health and reported fewer long-standing illnesses, disabilities, or infirmities (Fig. 2d left). This observation might indicate either superior health in the low neuroticism group or their propensity for a more positive health perception. Conversely, the high ERIS group engaged in more frequent suspect disease check-ups and medical follow-up examinations than the low neuroticism group (Fig. 2d right), suggesting that diligent health self-monitoring in the high ERIS individuals could lead to proactive health management, potentially contributing to their longer lifespan.

After evaluating the survival curves of different subgroups, our study next examined the associations between ERIS, neuroticism, and mortality in the top 5 high-fatality diseases (including ischemic heart disease, lung cancer, cerebrovascular disease, and breast cancer) and COVID-19. Employing Cox proportional hazards models adjusted for age, sex, and age-sex interaction, we identified a significant impact of ERIS on overall mortality (HR = 0.88, 95% CI: 0.87-0.89), particularly notable in ischemic heart disease(HR = 0.83, 95% CI: 0.79-0.86), lung cancer(HR = 0.79, 95% CI: 0.76-0.83), and COVID-19 (HR = 0.83, 95% CI: 0.77-0.89) (see Fig. 2e). In contrast, the neuroticism score exhibited a larger effect on the mortality associated with cerebrovascular disease compared to ERIS (see Fig. 2e). This pattern of associations was robust when using different confounds adjustments (such as body mass index, Townsend deprivation index, and qualification, refer to Supplementary Table 7). These findings indicate that the impacts of ERIS and neuroticism may vary across different causes of mortality.

### Two geometric gradients of neuroticism are associated with distinct disease and lifestyle characteristics

We further investigated the correlations between the ERIS and neuroticism with various diseases, including those with the highest global burden (see Methods). These diseases were categorized into two main groups: physical and mental illnesses. As shown in Fig. 3a-b, binary logistic regression analyses indicated that elevated neuroticism levels generally correspond to an increased prevalence of these diseases, except for stomach cancer, where the association was not significant (*P* > 0.05, FDR corrected). Conversely, higher ERIS scores were associated with a reduced prevalence of these diseases, with schizophrenia as an exception, showing a negative correlation with ERIS. To illustrate the specific impact of ERIS and neuroticism on disease susceptibility, we computed the relative differences in log odds ratios (log(OR_ERIS_) + log(OR_neuroticism_)) and displayed them as a gradient from left to right, showing a transition from ERIS’s protective effects to neuroticism’s increased susceptibility (Fig. 3a-b). Regarding physical illnesses, elevated ERIS levels are more specifically linked to road traffic injuries from motor vehicle (OR_ERIS_ = 0.85, 95% CI: 0.82-0.88), stomach cancer (OR_ERIS_ = 0.89, 95% CI: 0.84-0.95), diabetes (OR_ERIS_ = 0.79, 95% CI: 0.78-0.80), lung cancer (OR_ERIS_ = 0.83, 95% CI: 0.81-0.86), and COVID-19 (OR_ERIS_ = 0.83, 95% CI: 0.81-0.86), whereas neuroticism was more explanatory of headache disorders (OR_neuroticism_ = 1.33, 95% CI: 1.30-1.37), diseases of the oesophagus, stomach and duodenum (OSDD, OR_neuroticism_ = 1.29, 95% CI: 1.28-1.30), noninfective enteritis and colitis (OR_neuroticism_ = 1.24, 95% CI: 1.22-1.26), and gynaecological diseases (OR_neuroticism_ = 1.14, 95% CI: 1.13-1.16) (Fig. 3a). In terms of mental disorders, ERIS is more specifically associated with reduced tobacco use (OR_ERIS_ = 0.73, 95% CI: 0.71-0.74) and is weakly associated with anxiety disorders (OR_ERIS_ = 0.97, 95% CI: 0.95-0.98); neuroticism, however, shows a positive correlation with all types of mental illnesses (ORs_neuroticism_ range from 1.29 to 2.21, all *Ps* < 7.28×10^-35^) (Fig. 3b), aligning with previous studies supporting neuroticism as a potential general factor in psychopathology^1^.

**Fig. 3.**
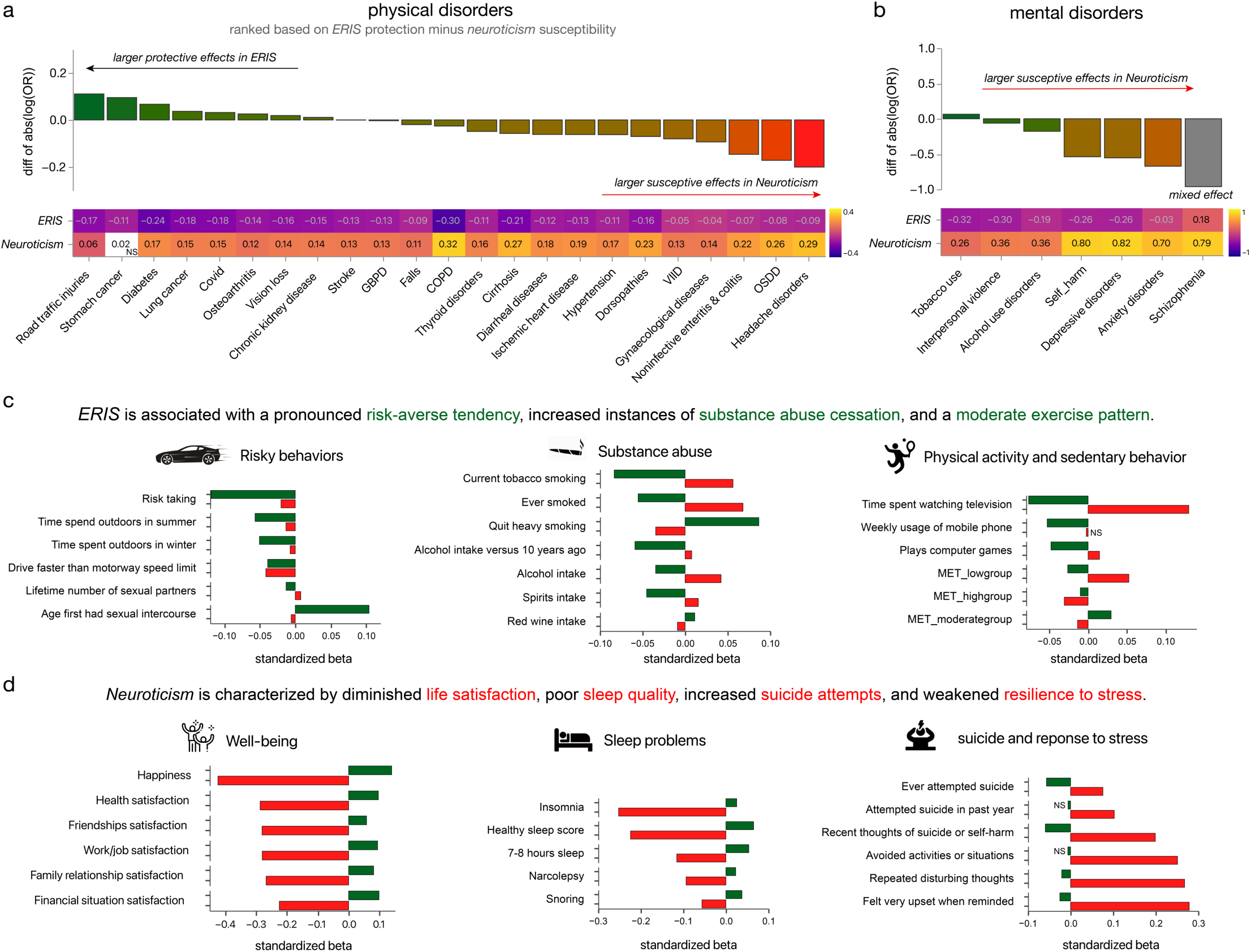
Correlation between neuroticism gradients and diseases/lifestyles. **(a-b)** Relationship between ERIS/neuroticism and the diagnosis of diseases classified according to ICD-10 **(a)** physical disorders; **(b)**mental disorders). The horizontal axis represents the 30 diseases selected for the study (as detailed in the Methods section). The lower part of the vertical axis shows the association coefficients between ERIS/neuroticism and these diseases, while the upper half of the vertical axis shows the difference in disease-related absolute log (odd ratio) values between ERIS and neuroticism. The associations of ERIS and neuroticism with different diseases are markedly distinct by log (odd ratio). **(c-d)** Relationships between ERIS/Neuroticism and the six lifestyle categories. The vertical axis shows the name of different lifestyles (see Methods), and the horizontal axis shows the standardized beta values of ERIS (green) and neuroticism (red) associated with the different lifestyles. Abbreviations: GBPD., disorders of gallbladder, biliary tract, and pancreas; COPD., chronic obstructive pulmonary disease; VIID., diseases of veins, lymphatic vessels and lymph nodes, not elsewhere classified; OSDD., diseases of oesophagus, stomach and duodenum. NS, not significant (*P* > 0.05 after FDR correction).

Regarding lifestyle characteristics, individuals with high ERIS personality traits typically show marked risk-averse tendencies, evidenced by a significant negative correlation with risk-taking behaviors (standardized *β* = - 0.12, *P* < 2.23×10^-308^). Interestingly, ERIS and neuroticism exhibit the same directional effects in this dimension; that is, individuals with low neuroticism levels are more inclined to engage in risk-taking (standardized *β* = -0.02, *P* = 5.55×10^-34^). The pattern of risk aversion among individuals with high ERIS traits extends to various risk-related behaviors, including speeding, engaging in certain sexual behaviors, and spending time outdoors during summer and winter (Fig. 3c). Notably, the variable ‘Age at first sexual intercourse’ shows that individuals within the lowest 5% of the ERIS range are approximately three times more likely to engage in early sexual activity (defined as age ≤15 years) compared to those in the highest 5%, for both male (7.66% for high ERIS vs. 22.02% for low ERIS) and female (5.28% for high ERIS vs. 15.19% for low ERIS). High ERIS are significantly linked with decreased substance use and more cessation behaviors. Remarkably, high ERIS and low neuroticism exhibit comparable effect sizes regarding the propensity to ‘Ever smoked’. However, ERIS shows significantly greater specificity for ‘Quit heavy smoking’ (Fig. 3c, ERIS: standardized *β* = 0.09, *P* = 5.74×10^-193^; neuroticism: standardized *β* = -0.03, *P* = 9.89×10^-34^). Interestingly, although high ERIS and low neuroticism similarly influence the overall ‘alcohol intake frequency’, individuals with high ERIS tend to consume less harmful alcohol types like spirits and more potentially beneficial types such as red wine (Fig. 3c). Additionally, individuals with high ERIS display significantly reduced ‘alcohol intake versus 10 years ago’, surpassing the effect size observed for neuroticism (Fig. 3c, ERIS: standardized *β* = -0.06, *P* = 3.69×10^-277^; neuroticism: standardized *β* = 0.01, *P* = 2.80×10^-6^). These results suggest that high ERIS may be particularly effective in prompting the cessation of unhealthy habits due to health concerns. In terms of physical activity and sedentary behavior (Fig. 3c), individuals with high ERIS predominantly engage in moderate-intensity exercises (‘MET_moderategroup’ refers to ‘MET minutes per week for moderate activity’; ERIS: standardized *β* = 0.029, *P* = 2.22×10^-63^; neuroticism: standardized *β* = -0.014, *P* =1.35×10^-14^) and generally avoid both high and low-intensity exercises. Conversely, those with low neuroticism exhibit a preference for high-intensity activities (Fig. 3c). Furthermore, high ERIS is associated with less use of mobile devices, playing computer games, and mobile phone usage (Fig. 3c). Lifestyle traits specifically linked to lower neuroticism are intuitive and widely corroborated by earlier research, showing a link with improved well-being and quality of life across various domains^27,28^. These domains include better life satisfaction, improved sleep quality, fewer suicidal attempts, and reduced adverse reactions to stress (Fig. 3d).

In addition, we examined the associations between different gradients of neuroticism with other domains. Notably, we found that individuals high in ERIS had similar positive associations with intelligence, educational attainment, socio-economic status and parental longevity as those low in neuroticism (see Supplementary Fig.2). In comparison to individuals with low neuroticism, those with high ERIS also demonstrated a dietary preference for cereals, cheese, and bread, while showing an aversion to unprocessed meats (see Supplementary Fig.2). Additionally, there was an increased preference for red wine (see below). This dietary pattern, characterized by a preference for cereals, fruit, fish, and a daily consumption of red wine, aligns closely with the traditional Mediterranean diet, renowned for its substantial health benefits^29^. Collectively, these results suggest that individuals with high ERIS exhibit more risk-averse and healthier lifestyle choices, potentially explaining their lower incidence of traffic accidents, COVID-19 conditions, and lifestyle-related chronic diseases, ultimately contributing to increased longevity.

### The neurostructural signatures of the geometric gradients of neuroticism

We next aimed to investigate the neuroanatomical basis of neuroticism gradients. Previous meta-analyses have failed to establish a consistent link between neuroticism and brain structures, such as gray matter volume^30,31^. Therefore, we aim to establish a stable relationship between neuroticism gradients and brain structures in a large sample, which will help provide deeper biological insights into the neuroanatomical basis of neuroticism.

Through a large-sample voxel-based morphometry (VBM) analysis of neuroticism in the UK Biobank (*n*=38,987), we identified distinct gradients of neuroticism attributable to mostly unique variations in gray matter volume across the entire brain. Fig. 4a illustrates voxels significantly associated with ERIS and overall neuroticism scores (Fig. 4a, voxel-level threshold *P* < 1×10^-3^ for ERIS and *P* < 0.05 for neuroticism, FDR corrected, cluster size > 100). We found a generally negative association of neuroticism with higher-order emotional brain regions, including the anterior cingulate cortex (ACC), medial prefrontal cortex, orbitofrontal cortex, anterior insula, and retrosplenial cortex, alongside a significant positive correlation with the left caudate. In contrast, inter-individual differences in ERIS demonstrated larger effects in associations with brain structure (Fig. 4b, with 15% of grey matter voxels showing absolute z-score values greater than 4, as opposed to only 1.5% for neuroticism), predominantly positive associations with the largest effect sizes observed in the cerebellum, medial thalamus, amygdala, hippocampus, parahippocampal gyrus, bed nucleus of the stria terminalis (BNST), and cortical areas such as the orbitofrontal cortex and insula. To enhance interpretability, we also used multiple brain atlases to perform ROI-based analyses (see Methods, Supplementary Table 8). Notably, these brain areas significantly overlap with established key nodes within the anxiety and fear circuits identified in animal models^32^. We next integrated the significant top 10% voxels related to ERIS and neuroticism into a Neurosynth functional annotation^33^ analysis (excluding the caudate due to its directionally inconsistent association). The results suggested that neuroticism is primarily linked to higher-order cognitive functions such as decision-making, strategy formulation, and monitoring (Fig. 4c), whereas ERIS aligns more closely with fundamental emotional functions like fear, anxiety, and risk assessment (Fig. 4d).

**Fig. 4.**
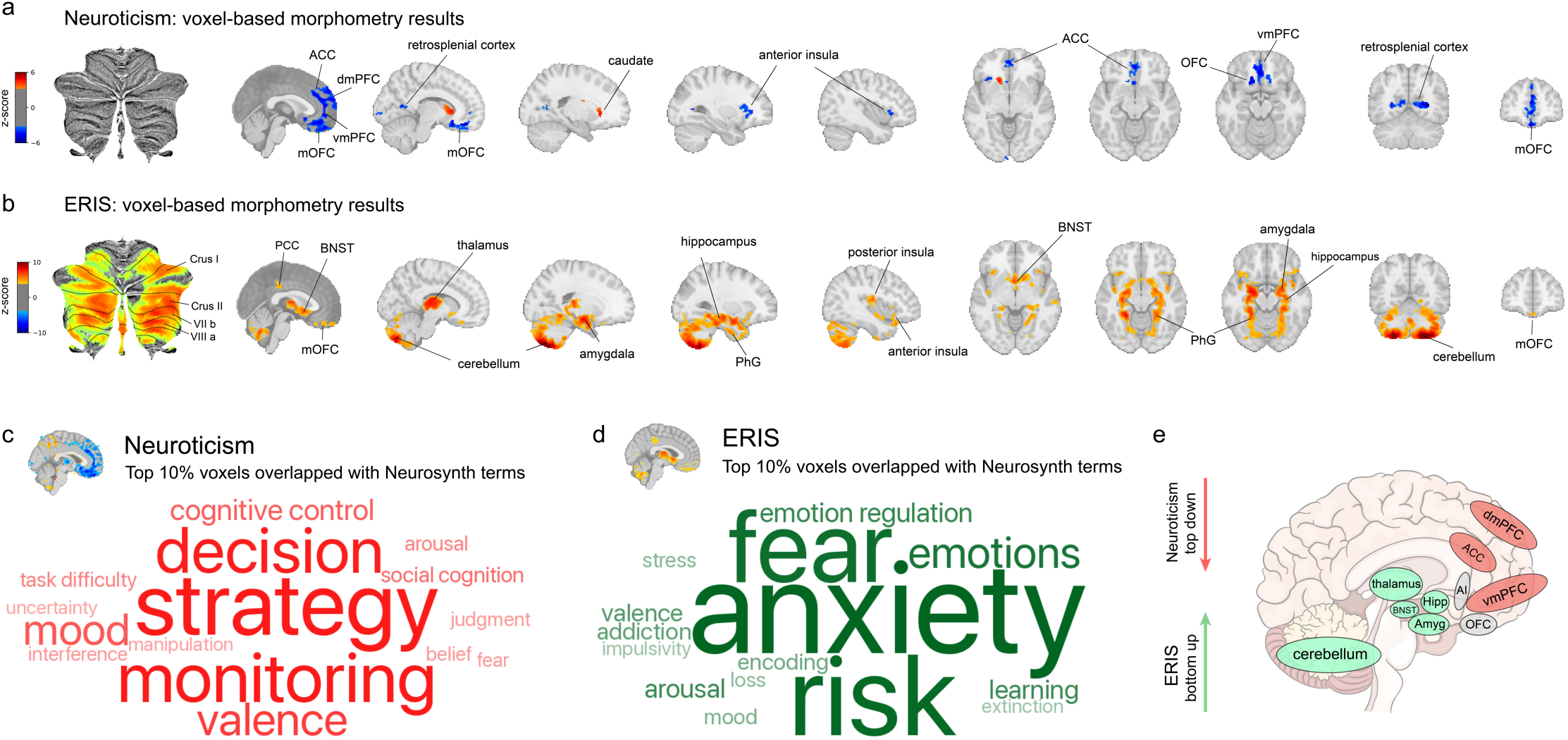
The neuroanatomical signatures of neuroticism gradients. **(a-b)** Voxel-wise regions significantly correlated with neuroticism **(a)** and ERIS **(b)** from VBM analysis. Neuroticism is primarily associated with grey matter volumes in higher-order brain regions, in particular the medial prefrontal cortex and the anterior cingulate cortex. Conversely, ERIS shows significant correlations with grey matter volume in more conserved areas, including the cerebellum, amygdala, hippocampus and thalamus. **(c-d)** Word cloud visualization of top 15 Neurosynth items related to neuroticism **(c)** and ERIS **(d)**, with font linearly corresponding to loading weights (see Methods). Neuroticism (red) is primarily associated with high-order cognitive functions such as decision-making, strategic formulation, and monitoring. ERIS (green) shows a specific association with the basic emotional responses, including fear, anxiety, and risk processing. **(e)** A schematic regarding the neurostructural signatures of neuroticism/ERIS is represented. Regions (red) linked to neuroticism are more specific in top-down cognitive modulation, whereas regions (green) related to ERIS are conservate, and more likely involved in the bottom-up process. The overlapped regions are shown in grey. Abbreviations: ACC., anterior cingulate cortex; dmpFC., dorsomedial prefrontal cortex; vmPFC., ventromedial prefrontal cortex; mOFC., medial orbitofrontal cortex; Crus I., Crus I cerebellum; Crus II., Crus II cerebellum; VIIIa, VIIIa cerebellum; VIIb., VIIb cerebellum; PCC., posterior cingulate cortex; BNST., bed nucleus of the stria terminalis; Phg., parahippocampal gyrus.

In summary, ERIS and neuroticism are linked to a distributed network of emotional circuits within the brain. Neuroticism primarily engages regions associated with higher-order emotion processing, indicating an involvement of top-down regulation mechanisms. Conversely, the neural correlates of ERIS are anchored in more anatomically conserved structures (see Fig. 4e, top 15 items are shown). These observations suggest a functional divergence between neuroticism, which may rely on cognitive modulation of emotions, and ERIS, which appears to depend on more fundamental, instinctual emotion responses.

### Genome-wide association studies (GWAS) of the geometric gradients

We conducted GWAS on geometric gradients in a cohort of 267,262 individuals from the UK Biobank. For the newly characterized phenotype ERIS, we identified 12 significant genomic risk loci (see Methods, Supplementary Table 9). The Manhattan plot for ERIS exhibited a *P* value association pattern distinctly different from that of neuroticism (Fig. 5a, top: ERIS; bottom: neuroticism). A review of the GWAS catalog^34^ revealed that 2 of the 12 loci were not associated with any neuroticism-related trait or psychiatric disorder (see Supplementary Table 9, chr7:10,990,005-11,136,052, chr16:7,215,912-7,237,717). Similar to phenotypic correlations, these loci were associated with risk-taking (9 loci), smoking (8 loci), and sex-related behaviors (13 loci), as detailed in supplementary data. The large sample size resulted in expected genomic inflation (ERIS: λ_GC_ = 1.29; Neuroticism: λ_GC_= 1.48) and linkage disequilibrium score regression^35^ (LDSC) intercepts close to 1 (ERIS: *intercept* = 1.023, *SE*= 0.0086; Neuroticism: *intercept* = 1.027, *SE* = 0.012), indicating that the signals are primarily attributable to true polygenicity rather than confounding biases. The LDSC SNP-based heritability (h^2^) for ERIS was estimated at 0.069 (*SE* = 0.0033) and for neuroticism at 0.116 (*SE* = 0.0060).

**Fig. 5.**
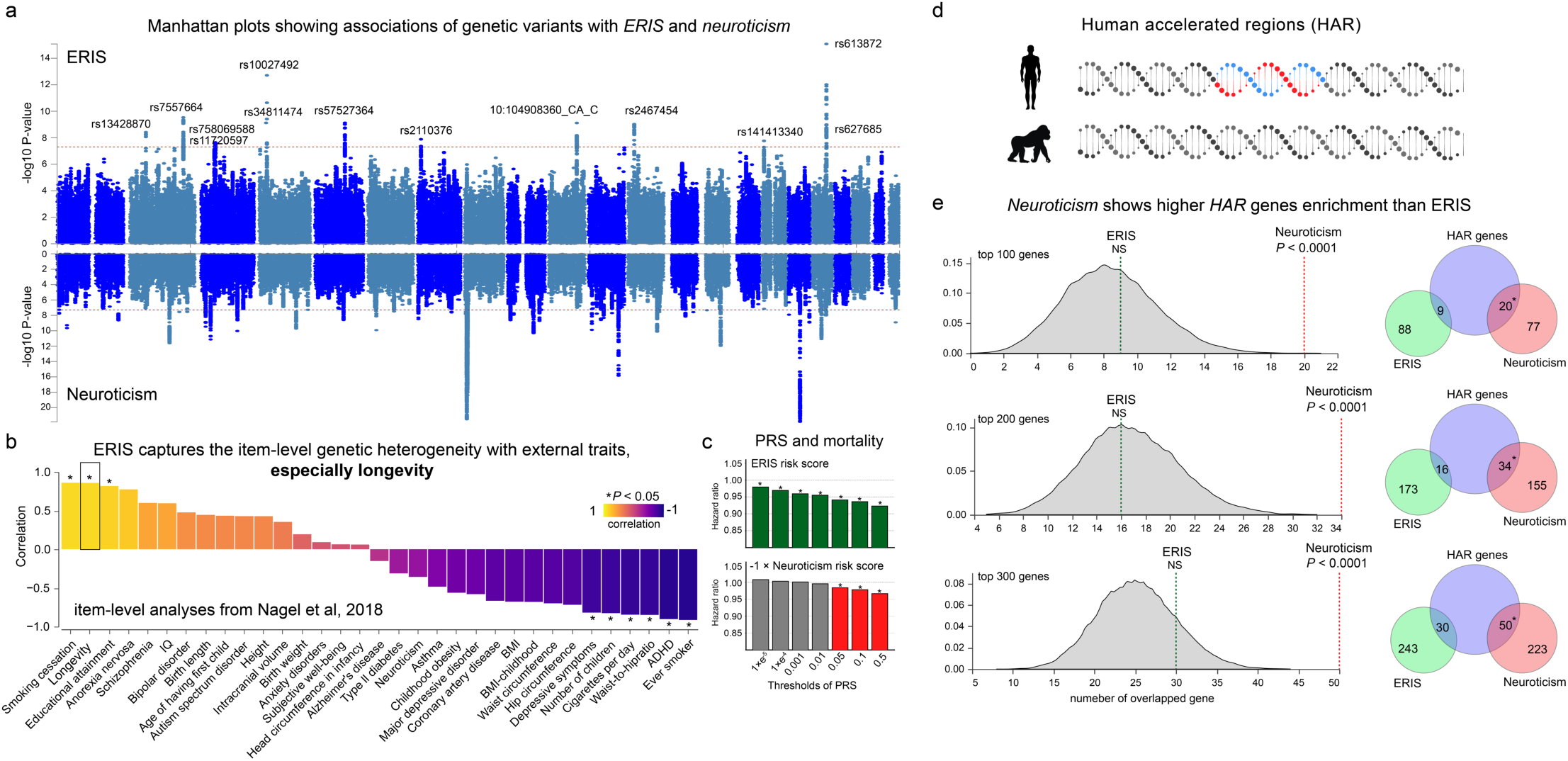
Genetic associations of neuroticism gradients. **(a)** Manhattan plot illustrating gene variants associated with ERIS and neuroticism. The genome-wide significance threshold (*P* < 5 × 10^-8^) is represented by a red line. The upper part of the plot highlights genomic loci associated with ERIS, while the lower part shows loci associated with neuroticism. **(b)** Bar chart showing the association between ERIS item weights and the item-level genetic correlations for 33 behavioral phenotypes derived from Nagel’s study^15^. Statistically significant correlations are marked with asterisks. The bars are ordered according to the size of the correlation coefficients. **(c)** Bar graph showing the relationship between different polygenic risk score (PRS) thresholds for ERIS/neuroticism and all-cause mortality. The x-axis represents different thresholds of the ERIS/Neuroticism PRS, while the y-axis represents hazard ratios from Cox proportional hazards regression models. **(d)** Schematic illustrating the analysis of genes associated with HARs, which represent genomic loci with accelerated divergence during human evolution. **(e)** Comparative analysis of gene expression levels within HARs for ERIS and neuroticism. The distribution plot on the left shows the distribution of HARs in randomly selected gene sets of 100, 200 and 300 genes. The vertical dashed lines indicate the number of genes significantly associated with ERIS (green) and Neuroticism (red). The Venn diagram on the right shows the frequency of occurrence of HARs in gene sets randomly associated with ERIS (green) or with Neuroticism (red).

Several studies have elucidated genetic heterogeneity within the structure of neuroticism^15,36^. Nagel et al.^15^ explored genetic correlations between 12 neuroticism items and 33 external traits using GWAS summary statistics independent of the UK Biobank. This exploration revealed that many items exhibited opposing effects. Building the result of Nagel et al, we investigated the association between the weights of ERIS (as depicted in Fig. 1d) and the heterogeneous genetic correlations across the 12 neuroticism items with external traits, aiming to pinpoint traits demonstrating genetic specificity with ERIS. Among the 33 external traits, the ERIS gradient showed significant positive correlations with smoking cessation, longevity, and educational attainment. Conversely, negative correlations were observed with ever smoking, attention deficit hyperactivity disorder, waist-hip ratio, daily cigarette consumption, and number of children. Remarkably, this analysis also confirmed a positive association between the ERIS and longevity (*r* = 0.86, *P* < 0.05, across 12 items) from a genomic perspective, independent of the death records from the UK Biobank cohort (see Fig. 5b). Subsequently, we calculated the polygenic risk scores (PRS) for ERIS and neuroticism and explored their associations with mortality. Utilizing Cox regression analysis (see Fig. 5c), we determined that a higher PRS for ERIS significantly predicts a reduced mortality rate across various PRS thresholds. While the PRS for neuroticism also predicts mortality, it does so with a lesser effect and without consistent impacts across different thresholds.

Inspired by our VBM results, we hypothesize that ERIS and neuroticism may exhibit varying degrees of evolutionary conservation, with neuroticism likely being more significantly expanded in recent human brain evolution. Consequently, we conducted an enrichment analysis that integrates GWAS results with genes associated with the human accelerated regions^37,38^ (HARs) (see Fig. 5d), previously identified to be highly expressed in the default mode network, such as the medial prefrontal cortex^38^, which showed associations with neuroticism. Using multi-marker analysis of genomic annotation (MAGMA) analysis^39^, facilitated by FUMA^40^, we identified 247 genes specifically associated with ERIS and 787 genes specifically associated with neuroticism, as well as 63 genes associated with both (*P* < 0.05, FDR corrected; see Supplementary Data). Permutation tests revealed that genes related to neuroticism, as opposed to ERIS, are significantly enriched in the HARs (neuroticism overlap: 127/787 genes, *P* < 1×10^-5^; ERIS overlap: 29/247 genes, *P* > 0.05). Results of gene-set analyses using MAGMA are presented in supplementary data. Selecting the top 100, 200, and 300 genes of ERIS and neuroticism yielded similar results (see Fig. 5e, the overlapped genes were removed).

Alternatively, selecting genes based on positional mapping in FUMA revealed that the results are robust (neuroticism overlap: 68/524 genes, *P* < 1×10^-5^; ERIS overlap: 12/149 genes, *P* > 0.05). These findings suggest that neuroticism, compared to ERIS, tend to have a higher rate of genetic evolutionary acceleration.

## Discussion

Our study establishes a multidisciplinary framework for personality research, utilizing a data-centric methodology to decipher latent population structures and their links to health outcomes, disease susceptibility, lifestyle factors, as well as neurobiological and genetic underpinnings. For neuroticism, which assesses an individual’s propensity for negative affect in response to adverse events, we identified two intrinsic gradients: 1) a general tendency toward response intensity, as indicated by the neuroticism total score; 2) the ERIS, delineating individuals’ inclination towards either adaptive response (such as worry or tension) or maladaptive response (such as irritability or mood instability) in moderately intense situations. We found that both a lower neuroticism score and a higher ERIS can predict longer lifespan, albeit through different mechanisms. Individuals with lower neuroticism likely benefit from improved well-being, better sleep, and reduced stress, decreasing their risk of mental and certain physical illnesses, especially gut and immune-related. In contrast, those with higher ERIS may achieve longevity through adaptive vigilance and risk avoidance, leading to fewer risky behaviors and substance abuse, better compliance with health screenings, and reduced susceptibility to chronic diseases like diabetes and cancer. Intriguingly, our study reveals that the ERIS dimension, compared to classic neuroticism, exhibits greater evolutionary conservation in anatomical and genetic aspects and has a stronger prediction power for future mortality.

Based on our findings, we propose an evolutionary hypothesis model of neuroticism, as illustrated in Fig. 6. Current human neuroticism encompasses two distinct dominant dimensions: high ERIS and low overall neuroticism, which are associated with optimal longevity and well-being, respectively. The former, likely an ancient trait, appears to be an inheritance from the shy-bold continuum^41^, observed across various species from fish to humans. This trait is linked to more conservative emotional circuits processing the fear and anxiety related to threats. In contrast, the latter dimension is a human-specific behavioral variation that has emerged more recently in response to ecological and cultural changes. This trait is characterized by higher-order emotional control circuits that refine strategies to manage and regulate stress responses, thereby enhancing happiness and overall life quality. Due to resource constraints, our ancestors and animals often engaged in trade-offs among various demands, such as those between survival and reproduction, or between starvation and predation. Across many species, the shy-bold (reactive-proactive) continuum^41–43^ has been consistently identified as a fundamental ‘animal personality’, wherein bolder individuals exhibit a higher propensity for risk-taking in reproduction and predation rather than prioritizing their survival and self-maintenance (bold individuals tend to reproduce more but survive less than shy ones). The ERIS dimension is likely intertwined with the shy-bold continuum. The positive extreme of the ERIS axis, which is typified by heightened emotional reactivity (e.g., anxiety), may encourage risk-averse behaviors^44^. Conversely, the negative extreme, characterized by emotional instability or distress, could amplify impulsiveness^44,45^. Our research underscores a pronounced association between ERIS and risk-taking behaviors, which surpasses that observed in overall neuroticism. This trend is strikingly apparent in the sexual domain, particularly the ‘age first had sexual intercourse’, where individuals in the lowest 5% ERIS bracket are approximately three times more likely to engage in early sexual activity compared to those in the highest 5%. Furthermore, we have found that the item-level specificity of ERIS is genetically linked to traits like ‘longevity’ (positive) and ‘number of children’ (negative), as evidenced in Fig. 5b. Therefore, we postulate that individuals with high ERIS scores, who typically adopt a cautious modern lifestyle, may embody an evolutionary adaptation of the shy personality (driven by an intrinsic survival instinct and a fear of mortality). In contrast, within the dimension of neuroticism, we postulate that lower levels of neuroticism have evolved into a predominant personality trait. This evolution facilitates the fulfilment of higher-order emotional needs (such as well-being), transcending the mere survival-related trade-offs as humanity has advanced. Modern humans are primarily faced with stressors related to daily life, such as academic pressure, social interactions, and workplace challenges, rather than threats to survival like predators. Individuals who are less affected by these modern stressors exhibit reduced stress responses, higher levels of happiness, life satisfaction, and sleep quality, all of which contribute to their overall physical and mental health. When considering longevity and well-being as integrated readouts, our findings suggest both high ERIS and low neuroticism as two different dominant strategies.

**Fig. 6.**
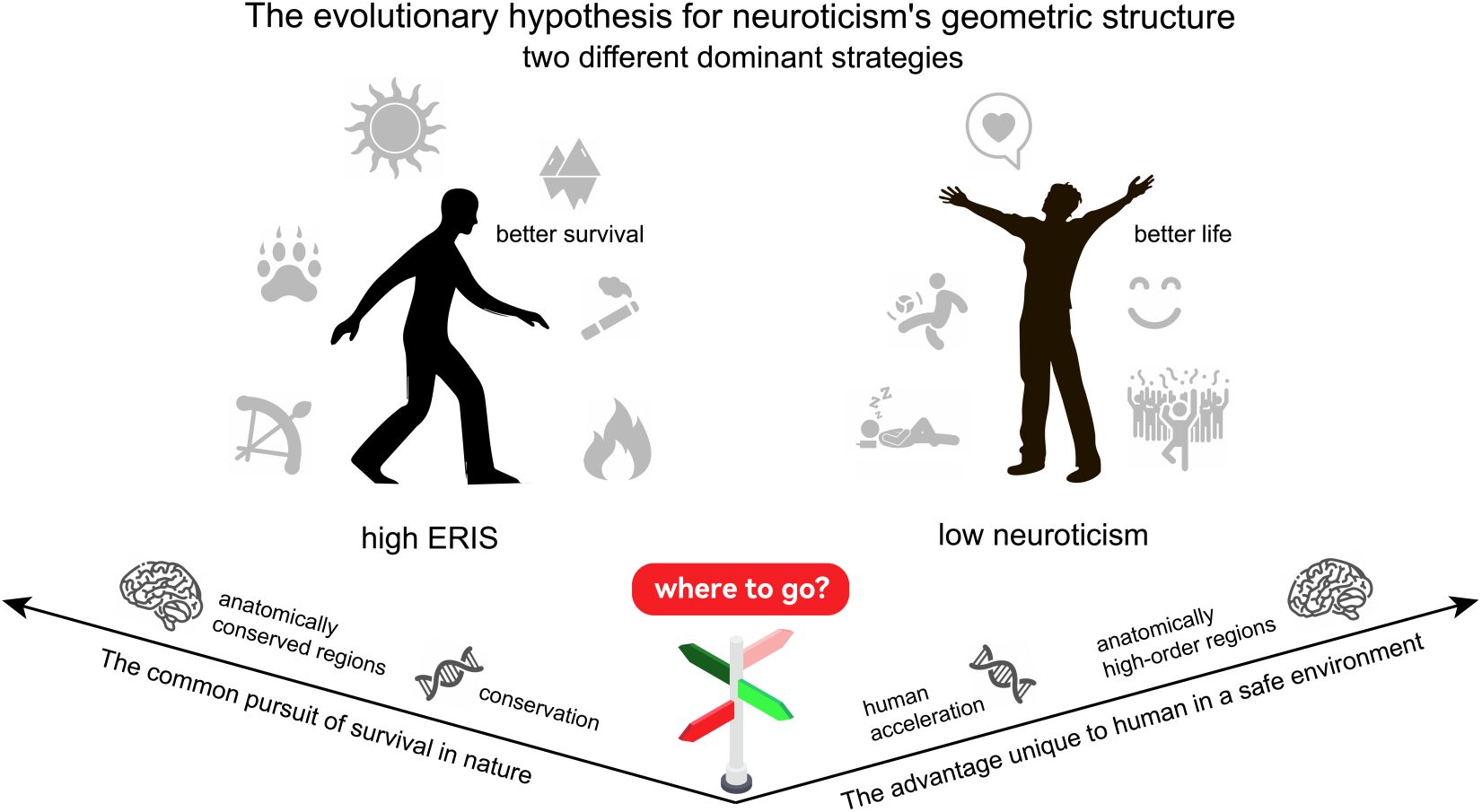
Evolutionary hypothesis of neuroticism’s geometric structure. This figure illustrates an evolutionary hypothesis model for the geometric structure of neuroticism, which outlines two dominant strategies inferred from the results of this study. The model argues that the geometric gradient of neuroticism encompasses two primary continuous spectra, within which high ERIS and low neuroticism emerge as two extreme groups, each endowed with distinct adaptive advantages. Individuals with high ERIS scores generally exhibit risk-averse behaviors that correlate with increased longevity. ERIS is not associated with HARs, but aligns with evolutionarily conserved brain regions primarily involved in primitive anxiety and fear circuits. In contrast, our analysis links neuroticism to HARs and higher-order brain regions, suggesting that neuroticism is a personality trait that evolved later, primarily to cope with stress and modulate stress responses. Individuals with lower levels of neuroticism often adopt a ‘pursuit of a better life’ strategy, which is associated with improved sleep, reduced stress responses, and increased well-being, reflecting a uniquely human strategy for achieving a fulfilling life.

Individuals with high neuroticism, when confronted with negative external stimuli, are often perceived as less effective in regulating negative emotions. Contemporary research^46,47^ frames emotion regulation as a dynamic interplay between cortical cognitive control systems, which orchestrate ‘top-down’ modulation, and subcortical systems, facilitating ‘bottom-up’ emotional generation. Our findings align with this perspective, indicating that the two geometric gradients of neuroticism correspond to distinct neuroanatomical bases along this hierarchy of emotional processing, as corroborated by Neurosynth results.

Specifically, individuals with high overall neuroticism tend to exhibit reduced grey matter volume in key cortical regions involved in the downregulation of negative emotions. These areas include the dorsomedial prefrontal cortex (dmPFC), ventromedial prefrontal cortex, ACC, anterior insula, and retrosplenial cortex. Particularly, the dmPFC plays a crucial role in cognitive processing like cognitive reappraisal, selective attention, and rumination, exerting top-down control over downstream emotional expression^48–50^. Similarly, the anterior insula^51,52^, rather than the posterior, is involved in emotion awareness, aligning with our results that link the anterior insula more with neuroticism, while ERIS does not show this specificity. Furthermore, the second gradient in the population, ERIS, exhibits distinct associations with evolutionarily conserved brain regions, including the cerebellum and limbic structures such as the thalamus, hippocampus, amygdala, and BNST. Echoing cues from Neurosynth and previous animal studies, these brain areas are integral nodes in anxiety and fear circuits. Recent research also increasingly acknowledges the significant role of the cerebellum^13,53–55^ in emotion processing. Genetically, neuroticism-associated genes are significantly enriched in human-accelerated genes, which are more expressed in higher-order cognitive networks. Considering above, we speculate that the individual differences in the overall neuroticism trait predominantly depend on higher-order capacity of ‘top-down’ regulation, while the ERIS trait is closely related to more conserved ‘bottom-up’ emotional reactivity.

Neuroticism is recognized as a fundamental domain of personality with substantial public health significance. Understanding its structure and origins may improve strategies for screening high risk population and providing effective prevention methods. Beyond the classic neuroticism score, we show distinct neuroticism geometric gradients correlate with differing susceptibilities to diseases and mortality rates, allowing for the stratification of populations into various risk categories, such as those with high neuroticism or low ERIS. Previous studies^46^ indicate that cognitive reappraisal is more effective in regulating top-down than bottom-up generated emotions. Considering the distinct neuroanatomical correlates of overall neuroticism and ERIS, our findings imply the potential for varied intervention strategies. For example, it warrants investigation whether individuals with low ERIS would be more responsive to bottom-up approaches^47^ such as experiential emotion regulation or physical exercise, while those with elevated neuroticism levels may benefit more from cognitive behavioral therapy.

In our work, we developed a subject-level similarity network based on neuroticism item scores and employed diffusion map embedding to identify key neuroticism gradients. This concept and approach, contrasting with classic psychometrics reliant on closely positive item correlations, is designed to uncover the main dimensions of neuroticism’s population structure with minimal assumptions about weight directionality. For instance, ERIS is characterized by both positive and negative weightings, a concept not typically endorsed in factor analysis. Our findings suggest that such a personality trait with mixed weights has significant implications and strengthens correlations with behavioral and biological phenotypes. Furthermore, these geometric gradients of neuroticism are consistently observable across five independent cohorts.

Several previous studies have suggested the heterogeneity within the neuroticism domain. Nagel et al.^15^ illuminated the genetic correlations between item-level neuroticism and 33 distinct psychological traits, a result used in our work to address the genetic specificity of ERIS. Gale et al. employed a bifactor model to identify two specific neuroticism factors^9^: anxiety/tension and worry/vulnerability, which mirror the positive part in ERIS (see Supplementary Fig.3). This study, along with several subsequent work^16^, found positive correlations of these factors with affluence, intelligence, health, and longevity from genetic or phenotypic standpoints. In our supplementary notes, we discuss the relationship between ERIS and the findings from previous studies. Extending to existing studies, our study strives to provide an integrated framework for elucidating the inherent structure of neuroticism, incorporating insights from psychology, data science, epidemiology, neuroscience, genetics, and evolution.

Several limitations in our study are noteworthy. The UK National Statistics Office reports that from 2018 to 2020, the average death age was 82.3 years for men and 85.8 years for women, whereas UK Biobank participants with death certificates had average death ages of 71.1 and 70.8 years, respectively. This discrepancy suggests limitations in our study due to the duration of longitudinal tracking. Future research with extended follow-up periods could yield deeper insights into neuroticism’s link with all-cause and specific disease mortalities. Our study used two different scales to examine neuroticism’s geometric structure across diverse, multi-centered cohorts. Notably, unlike the EPQ-N, the FFI-N does not measure mood instability directly. Future research should include ERIS items and investigate more specific neuroticism measurement scales. While our study focused on mortality, future studies should broaden the scope to include diverse health outcomes and employ multi-wave cohort designs to clarify neuroticism’s relationship with disease. It would also be beneficial to investigate interventions, such as lifestyle modifications, that could influence these associations.

## Method

### Neuroticism measurements

Neuroticism is defined as a relatively stable trait, marked by an inclination to experience a range of negative emotions. This spectrum includes, but is not limited to anxiety, depression, irritability, and emotional instability^2^. Although the concept of neuroticism is widely acknowledged, there are various versions of measurement questionnaires available. Among them, the FFI-N and the EPQ-N are widely recognized and utilized internationally for evaluating neuroticism^56,57^. Overall, the FFI-N and the EPQ-N cover similar domains. However, the EPQ-N offers a more direct measurement of mood swings, an aspect not explicitly addressed in the FFI-N. A recent study highlights the pivotal role of negative emotions in the construct of neuroticism, an aspect where the objectives of both questionnaires converge^58^.

#### FFI-N (HCP, HCP-D, CN-U, and CN-A datasets)

The FFI-N, developed by Paul T. Costa and Robert R. McCrae, consists of 12 items designed to assess neuroticism on a Likert 5-point scale, with higher scores indicating greater levels of neuroticism^59^. Extensive past research has validated the high reliability and validity of the English version, establishing it as a trustworthy tool for neuroticism assessment^60,61^. This version is used in the HCP and HCP-D datasets, with specific questionnaire items detailed below. For the Chinese version, we calculated the Cronbach’s alpha coefficient to verify its reliability, as elaborated in the supplementary notes. The Chinese version was administered in the CN-A and CN-U samples, with details of the questionnaire items available in the supplementary notes. To provide a comprehensive overview of the scale’s application across these datasets, Supplementary Fig.4-7 respectively display the frequency of each neuroticism item within the four datasets.

The English version utilized in the HCP and HCP-D datasets is:

1. I am not a worrier (abbreviated as **Worried**)
2. I often feel inferior to others (abbreviated as **Inferior**)
3. When I’m under a great deal of stress, sometimes I feel like I’m going to pieces (abbreviated as **Stress**)
4. I rarely feel lonely or blue (abbreviated as **Blue**)
5. I often feel tense and jittery (abbreviated as **Tense**)
6. Sometimes I feel completely worthless (abbreviated as **Worthless**)
7. I rarely feel fearful or anxious (abbreviated as **Anxious**)
8. I often get angry at the way people treat me (abbreviated as **Irritability**)
9. Too often, when things go wrong, I get discouraged and feel like giving up (abbreviated as **Discourage**)
10. I am seldom sad or depressed (abbreviated as **Depressed**)
11. I often feel helpless and want someone else to solve my problems (abbreviated as **Helpless**)
12. At times I have been so ashamed, I just wanted to hide (abbreviated as **Ashamed**)

#### EPQ-N (UK Biobank dataset)

The EPQ-N, conceived by British psychologist Hans Eysenck, encompasses 12 questions with response choices of ‘Yes’, ‘No’, ‘Don’t know’, and ‘Prefer not to answer’^21^. In our study, responses marked as ‘Don’t know’ and ‘Prefer not to answer’ were treated as missing data. This scale has been validated for its reliability and applicability across different cultural contexts. We visualized the item score frequencies of EPQ-N among UK Biobank participants in Supplementary Fig.8.

The specific questions of the EPQ-N are:

1. Does your mood often go up and down? (abbreviated as **Moodswing,** Field ID:1920)
2. Do you ever feel ‘just miserable’ for no reason? (abbreviated as **Miserable,** Field ID:1930)
3. Are you an irritable person? (abbreviated as **Irritability,** Field ID:1940)
4. Are your feelings easily hurt? (abbreviated as **Sensitivity,** Field ID:1950)
5. Do you often feel ‘fed-up’? (abbreviated as **Fed up,** Field ID:1960)
6. Would you call yourself a nervous person? (abbreviated as **Nervous,** Field ID:1970)
7. Are you a worrier? (abbreviated as **Worrier,** Field ID 1980)
8. Would you call yourself tense or ‘highly strung’? (abbreviated as **Tense,** Field ID:1990)
9. Do you worry too long after an embarrassing experience? (abbreviated as **Worry,** Field ID:2000)
10. Do you suffer from ‘nerves’? (abbreviated as **Nerves,** Field ID:2010)
11. Do you often feel lonely? (abbreviated as **Loneliness,** Field ID:2020)
12. Are you often troubled by feelings of guilt? (abbreviated as **Guilty,** Field ID:2030)

### The subjective categorization of the neuroticism items

Neuroticism is defined as a trait characterized by a predisposition towards negative emotional reactions, particularly in response to external stressors^62^. To improve clarity and enhance interpretation, we have subjectively divided the questionnaire items of the Five-Factor Inventory-Neuroticism (FFI-N) and Eysenck Personality Questionnaire-Neuroticism (EPQ-N) into two distinct domains: emotional reactivity and emotional instability/distress.

In our classification, Emotional Reactivity encompasses relatively common and basic negative affectivity that is typically triggered by adverse situations and stress, such as worry, anxiety, tension, and feelings of depression^63–65^. In contrast, Emotional Instability/Distress includes emotional variability and perceptible internal distress and turmoil, often linked with social contexts and involving more complex cognitive or emotional responses, such as feelings of shame and inferiority^66,67^. It is important to emphasize that this categorization, while subjective, is specifically designed to facilitate the interpretation and visualization of data (as illustrated in Fig. 1a) and does not influence the statistical outcomes of our analysis, since it was not incorporated into the data processing stage.

Specifically, we categorized the following items under the Emotional Reactivity domain due to their measurement of fundamental emotional reactions: from FFI-N, ‘Worried’, ‘Anxious’, ‘Depressed’, ‘Blue’, ‘Tense’, and ‘Stress’; and from EPQ-N, ‘Worrier’, ‘Worry’, ‘Sensitivity’, ‘Nervous’, and ‘Nerves’. Conversely, items that reflect a greater degree of social attributes and internal distress were assigned to the Emotional Instability/Distress domain: from FFI-N, ‘Inferior’, ‘Irritability’, ‘Discourage’, ‘Worthless’, ‘Helpless’, and ‘Ashamed’; and from EPQ-N, ‘Loneliness’, ‘Guilty’, ‘Irritability’, ‘Fed Up’, and ‘Miserable’. The ‘Moodswing’ item from EPQ-N, which directly assesses emotional variability, is also categorized under this domain.

Ambiguously, both the FFI-N item ‘I rarely feel lonely or blue’ and the EPQ-N item ‘Do you often feel lonely?’ include the concept of loneliness. We consider loneliness to be a form of emotional distress stemming from unmet social needs. However, the inclusion of ‘blue’ in the FFI-N item, closely related to the concept of being depressed, leads to its classification under emotional reactivity. On the other hand, ‘fed up’ and ‘miserable’, although not necessarily related to social attributes, are considered forms of distress and are thus included under emotional instability/distress.

### Participants

#### UK Biobank

In our research, we harnessed the extensive data resources of the UK Biobank, registered under application number 85139^68^. This comprehensive database includes a wide range of data, encompassing phenotypic, imaging, and genetic information^69,70^. Our study primarily focused on data from the initial registration phase (2006-2010) and the third visit phase (2014-2020), offering a wealth of individual participant information.

During the recruitment period (2006–2010), the UK Biobank conducted a thorough integration of diagnostic and evaluative methodologies. This approach involved detailed demographic surveys, evaluations of neuroticism, mental health, lifestyle questionnaires^71^. Additionally, blood samples were gathered for genome-wide genotyping, enabling in-depth biochemical analyses. To assess neuroticism, the UK Biobank used the 12-item EPQ-N and included 401,574 individuals (215,585 females, age: 56.41 ± 8.07 years old).

In the third visit phase (2014–2020) of the UK Biobank, neuroticism follow-up assessments were conducted for 35291 participants (17717 females, age: 54.92 ± 8.07 years old)^68^. Concurrently, structural brain imaging was carried out at imaging centers in Manchester, Reading, and Newcastle^68,70^. The neuroticism data from this phase were mainly used to perform VBM analysis.

Beyond these primary data collection periods, our study also incorporated additional longitudinal data, including National Health Service (NHS) medical records, which covered ICD-10 classifications and COVID-19 disease and mortality records^72,73^. This supplementary data facilitated our exploration of the relationships between neuroticism gradients, disease, and life expectancy.

The UK Biobank rigorously adhered to ethical standards and received approval from the North West Multi-centre Research Ethics Committee. Consistent with established research ethics practices, comprehensive informed consent was meticulously obtained from all participants. This protocol guaranteed the integrity and ethical compliance of our research, maintaining the highest standards of research ethics and safeguarding participant privacy.

#### HCP

Our study incorporated personality data from the HCP S1200 young adult subjects (n=1198; 650 females, age: 28.84 ± 3.68 years old)^74^. The HCP implemented the NEO Five-Factor Inventory (FFI), a comprehensive 60-item questionnaire tailored to evaluate the five-factor model of personality^59^. This instrument encompassed five dimensions: neuroticism, agreeableness, openness, conscientiousness, and extraversion^60^. Participants provided their responses on a 5-point Likert scale, varying from strongly disagree to strongly agree^75^. The FFI has been extensively validated in the United States and several other countries, confirming its reliability and applicability across diverse populations^76^. We explored the relationships between different gradients of neuroticism and FFI traits (see Supplement Fig. 10-11). Adjusted for age and sex, the first neuroticism gradient predominantly exhibited high correlations with neuroticism (partial *r* = 0.99, *P* < 2.2×10^-308^), and significant correlations with conscientiousness (partial *r* = -0.41, *P* = 1.06×10^-48^), extraversion(partial *r* = -0.34, *P* = 3.01×10^-34^) and agreeableness(partial *r* = -0.32, *P* = 5.25×10^-30^), while the ERIS showed positive correlations with conscientiousness (partial *r* = 0.20, *P* = 9.08×10^-12^), agreeableness (partial *r* = 0.14, *P* = 9.75× 10^-7^) and extraversion (partial *r* = 0.09, *P* = 1.14×10^-3^).

The data for this study, accessible via the HCP website (www.humanconnectome.org), complied with all pertinent guidelines and regulations and received approval from the Washington University Institutional Review Board. Accessing these data required adherence to the specific data use terms set by the HCP, which include protocols for managing both open access and restricted data. The latter category involves sensitive information such as exact age. This research was conducted in compliance with the HCP restricted data use terms, as per point 6 of the HCP data use document, this institution does not require a separate or individual ethics committee submission and/or approval.

#### HCP-D

We analyzed the Human Connectome Project Development (HCP-D) 2.0 release data^20^. This resource encompasses a developmental spectrum, focusing on individuals from 5 to 21 years of age, and is hosted across four distinguished U.S. sites: Harvard University, University of California, Los Angeles, University of Minnesota, and Washington University in St. Louis. For our study, we included personality questionnaire data from 229 participants (125 females; age: 19.00 ± 1.83 years). For the assessment of personality traits, the HCP-D adopted the same methodology as the HCP, utilizing the FFI questionnaire^59^. Within this framework, our analysis was narrowed down to data from 229 participants, comprising 104 males and 125 females. The age range of these participants was 16-21 years, as the personality assessment was conducted only on individuals over the age of 16. Similar to the HCP, the associations between different gradients of neuroticism and FFI personality dimensions were examined (see Supplementary Fig.9-10). Adjusted for age and sex, the first gradient of neuroticism was mainly characterized by correlations with traits such as neuroticism (partial *r* = 0.99, *P* = 5.48×10^-263^), openness(partial *r* = 0.17, *P* = 0.00975), agreeableness (partial *r* = -0.27, *P* = 3.80×10^-5^), conscientiousness (partial *r* = -0.38, *P* = 2.30×10^-9^) and extraversion (partial *r* = -0.38, *P* = 2.09×10^-9^). In contrast, the ERIS was positively associated with agreeableness (partial *r* = 0.25, *P* = 2.00×10^-4^) and conscientiousness (partial *r* = 0.33, *P* = 4.60×10^-7^).

The study was meticulously aligned with all relevant ethical guidelines and regulations. Informed consent was obtained from all participants, and the study received approvals from the Institutional Review Board at each participating site. This approach ensured the integrity and ethical compliance of the research process, reflecting a commitment to upholding the highest standards of research ethics and methodology.

#### CN-U and CN-A

We included two independent cohorts from mainland China: a group of CN-U and a group of CN-A. The university student cohort (CN-U) included 612 young adults (318 females, age: 20.12 ± 1.08 years old) from Central South University, Hunan, China. The adolescent cohort (CN-A) comprised 532 teenagers (253 females, age: 16.35 ± 0.92 years old) from a public middle school in Hunan, China. For the measurement of neuroticism in these two mainland Chinese cohorts, the FFI-N was utilized. It is noteworthy that for these participants, the focus was solely on the neuroticism dimension of the five-factor model of personality, without assessments of the other personality traits.

In both groups, the process of obtaining informed consent was conducted with adherence to ethical standards. Consent was directly acquired from participants or, for minors, from their guardians. The entire research protocol, encompassing the procedures for consent and data collection, was reviewed, and approved by the Ethics Committee of the School of Psychology at Capital Normal University.

### Low-dimensional Embeddings of Neuroticism

To quantitatively evaluate the heterogeneous subdimensions of neuroticism in individuals, we computed the primary and secondary embedding scores based on a between-subject similarity network. The definition of network similarity hinges on the levels of neuroticism sub-scales across different individuals. Two individuals can have identical overall neuroticism scores yet exhibit minimal similarity due to differing sub-scale profiles. Specifically, for n subjects, we first constructed a between-subject distance network *G* (*n*×*n*). In the context of a multi-item neuroticism questionnaire, the element *G_ij_* located in the i_th_ row and j_th_ column of matrix *G* quantifies the Euclidean distance between the item score vectors of subjects i and j. Subsequently, we identified the maximum value in *G*, denoted as G_max_. For each element *G_ij_*, the between-subject similarity network was derived by calculating (*G_max_* – *G_ij_*) / *G_max_*, yielding values ranging from 1 (maximum similarity) to 0 (minimum similarity). We employed diffusion map embedding^77^, a non-linear dimensionality reduction algorithm, to extract low-dimensional gradients of neuroticism, utilizing the BrainSpace toolbox^78^ (version 0.1.10). This method, known for its robustness against noise perturbations compared to algorithms like principal component analyses, is adept at identifying low-dimensional manifolds. The diffusion map embedding algorithm operates based on two key hyperparameters: *α* and *t*. The *α* parameter modulates the impact of the density of data points on the manifold (with *α* = 0 indicating maximum influence and *α* = 1 signifying no influence), while *t* determines the scale of the eigenvalues of the diffusion operator. In our analysis, we adhered to previously established values, setting *α* at 0.5 and *t* at 0. This configuration is chosen to preserve the global relationships among data points in the embedded space.

For the HCP, HCP-D, CN-U, CN-A, and UK Biobank datasets (including 0.0 and 2.0 time points), we computed individual gradient scores following the above pipeline. In the case of the UK Biobank (0.0 time point) dataset, due to its large size, we adopted a strategy to reduce computational memory requirements. Specifically, we randomly selected 50,000 participants for the calculation of the gradient scores. To extrapolate these scores to the entire cohort, we constructed regression models based on k-nearest neighbors (K = 50) using the gradient scores of these 50,000 individuals for different neuroticism gradients. This model was then applied to all participants. This process was repeated 100 times, yielding highly stable gradient scores across individuals. We determined the item-level weight contributions by correlating each individual’s gradient scores with their respective original item scores. Our study unveils the ERIS gradient as a stable phenotype with item-specificity. In the case of the FFI-N, item-level weights were compared across four independent datasets: HCP, HCP-D, CN-U, and CN-A. For the EPQ-N, item-level weights were compared across two different time points of the UK Biobank dataset, revealing remarkable consistency.

We next examined whether there was a significant correlation between the ERIS gradients derived from the FFI-N and the EPQ-N. Acknowledging the inherent differences between these scales, we employed a hybrid approach of quantitative and qualitative methods for item matching, aligning FFI-N items with those from the EPQ-N. Quantitatively, we utilized the ‘all-MiniLM-L6-v2’ model from the SentenceTransformers Python package^25^ to convert questionnaire items into embeddings. This facilitated the calculation of similarity indices between items from different questionnaires using cosine similarity. Based on the degree of similarity, pairs of items from the two scales were identified. These proposed pairs underwent two experts review for validation, resulting in the confirmation of eight matched pairs (see Supplementary Table 10). Following this, we transformed the FFI-N item weights from the HCP, HCP-D, CN-U, and CN-A datasets to align with those of the EPQ-N. This transformation facilitated a comparison of the item weights in FFI-N datasets and the EPQ-N UK Biobank dataset (as shown in Fig. 1g).

### Mortality Prediction and Cox Analysis

In the UK Biobank dataset, participants’ death certificates were sourced from the archives of the UK’s NHS. The precise date of death of participants was provided by the UK’s National Death Registry. This registry integrates data from NHS Digital, overseeing England and Wales, and the NHS Central Register, responsible for Scotland^79^. All records adhere strictly to the International Classification of Diseases, Tenth Revision (ICD-10), a WHO-developed system for consistent global disease classification^80^. Our study encompassed 36,366 death cases recorded between July 7, 2007, and November 12, 2021.

We examined the relationship between varying gradients of neuroticism and mortality risk, with a particular interest in comparing subgroups exhibiting minimal neuroticism levels. Within the UK Biobank cohort, we categorized the population into four distinct, non-overlapping subgroups based on gender and age: low neuroticism, high neuroticism, low ERIS, and high ERIS. Initially, we set the sample count for each subgroup starting with a neuroticism score of zero. Subsequently, we ranked the participants to categorize them into low neuroticism (with a sum score of zero), high neuroticism, low ERIS, and high ERIS groups. We then eliminated duplicates across these groups and equalized the group sizes by randomly downsizing the three larger groups to match the size of the smallest group, maintaining stringent control over age and gender variables, which are known to be closely associated with mortality outcomes (each group n=44,399). We then performed Kaplan-Meier survival analysis to produce survival curves for each subgroup over an approximate duration of 180 months, as depicted in Fig. 2b^81^. This analysis facilitated the assessment of cumulative survival probabilities at various intervals. For cases with missing data or unrecorded deaths, November 12, 2021, was used as a cutoff date. We applied the log-rank test to compare survival distributions across different subgroups^82^. Considering that random factors in population stratification could slightly influence the outcomes, we created 101 different grouping scenarios and reported a median effect size between the high ERIS and low neuroticism groups; furthermore, gender-specific log-rank tests were conducted to demonstrate significant differences between these groups among both males and females.

We subsequently focused on specific causes of death within the UK Biobank records, adhering to the ICD-10 system and classified under code 40000 and 40001. Due to participant number constraints, our analysis concentrated on the five leading causes of death and COVID-19 (see Supplementary Table 11). To evaluate time-dependent risks throughout the study, we employed the Cox proportional hazards regression model, a well-established semi-parametric method in survival analysis^83^. This model calculates the hazard ratio to quantify the change in death risk associated with each unit increase in the neuroticism gradient. Survival times were determined in months from the date each participant joined the study at the evaluation center until their death or the cut-off in November 2021. In our analyses, age and gender served as primary covariates, with additional strategic covariates included to broaden the robustness and scope of our study (detailed in the Supplementary Table 7).

Given our finding of a higher survival rate among individuals with high ERIS compared to those low in neuroticism, we further explored possible explanations by examining self-rated health, suspected medical examination, and post-treatment follow-up examination in two groups defined by high ERIS and low neuroticism (see Supplementary Table 12). The self-report health condition of the participants involved two questions: i) ‘In general, how would you rate your overall health?’ with responses categorized as ‘Poor’, ‘Fair’, ‘Good’, or ‘Excellent’ (Field ID: 2178); ii) ‘Do you have any long-standing illness, disability, or infirmity?’ with binary response options of ‘Yes’ or ‘No’ (Field ID: 2188). Furthermore, we examined all medical observations and evaluation for suspected diseases and conditions (ICD-10 code: Z03). For patients with hospitalization records, we examined all instances of post-treatment follow-up (ICD-10 code: Z08 and Z09). We employed the Chi-squared test to assess differences between groups characterized by high ERIS scores and low neuroticism scores^84^.

### Disease

Participants’ medical and diagnostic histories were extracted from the comprehensive medical archives of the NHS. Given the disparity in data coverage within the UK Biobank cohort, where primary care clinical records cover approximately 45% of the cohort compared to the more comprehensive 83% coverage by hospital inpatient records, our study primarily utilized the latter source^85^. This choice was made to ensure the robustness and completeness of the data analyzed, enhancing the reliability of our findings. These inpatient records were meticulously categorized using the ICD-10, focusing on diagnoses requiring hospital admission (as denoted by ID 41270 in the UK Biobank database).

Our investigation aimed to elucidate the relationship between different levels of neuroticism and specific disease risks, with a particular focus on the top 33 diseases contributing to the highest global burden of disease in adults aged 25-74 in 2019^86^. Maternal conditions were excluded from our analysis due to the scope of the study. A detailed process of adjustment and merging was applied to these conditions, tailored to the characteristics of our UK Biobank dataset, resulting in a refined list of 30 disease clusters. For example, conditions reported in The Lancet paper as ‘low back pain’, ‘neck pain’ and ‘other musculoskeletal’ were merged into ‘dorsopathies’, ‘hypertensive heart disease’ was subsumed under ‘hypertensive diseases’, and ‘road traffic injuries’ were classified as ‘motor vehicle road traffic injuries’. Tobacco use was also included in the analysis. In addition, conditions with fewer than 1,000 reported cases in the UK Biobank were excluded from the study. Conditions were divided into mental and physical disorders, with physical disorders including ‘motor vehicle road traffic injuries’ (V20-V79), ‘diabetes’ (E10-E14), ‘headache disorders’ (G43-G44), ‘diseases of oesophagus, stomach and duodenum’ (abbreviation as OSDD, K20-K31), ‘lung cancer’ (C34), ‘stomach cancer’ (C16) and others. Mental disorders included ‘depressive disorders’ (F32-F33), ‘schizophrenia’ (F20), ‘anxiety disorders’ (F40-F41), ‘alcohol use disorders’ (F10), ‘tobacco use’ (F17) and others. A comprehensive list of the selected disorders and the associated ICD codes can be found in the Supplementary table 13.

We applied a generalized linear model (GLM) to assess the relationship between neuroticism gradients and disease occurrence^83^. In this analysis, the dependent variable was the presence of disease (binary: Yes or No), while the independent variables included phenotype of interest (such as ERIS or neuroticism), sex, age, and the interaction between sex and age. The GLM analysis was conducted using the Python package statsmodels^87^.

### Lifestyle

Lifestyle choices, as manifestations of underlying psychological and behavioral predispositions, offer valuable insights into health outcomes associated with neuroticism gradients^2,88,89^. In this study, leveraging the expansive dataset from the UK Biobank, we focused on six lifestyle domains indicative of these choices: risk-taking behaviors, smoking habits, physical exercise, life satisfaction, sleep patterns, and responses to suicide/stress. Additionally, our supplementary results include an examination of dietary habits, the Townsend deprivation index, and other personal information (see Supplementary Fig.4). Data pertaining to lifestyle were primarily obtained using Category 100050 from the UK Biobank, with specific behaviors pinpointed through diverse Field IDs (see Supplementary Table 14). Responses classified as ‘Don’t know’ or ‘Prefer not to answer’ were systematically excluded.

#### Risk-taking behaviors

Neuroticism, characterized by an enhanced sensitivity to uncertainty and risk, displays an inconsistent relationship with risk-taking behaviors, as demonstrated by previous studies^90,91^. In our study, we included a set of risk-taking indicators: including self-reported ‘Risk taking’ (Field ID: 2040), ‘Drive faster than motorway speed limit’ (Field ID: 1100), and propensity for sexual behavior (‘Age first had sexual intercourse’, Field ID: 2139; ‘Lifetime number of sexual partners’, Field ID: 2149). Furthermore, we indirectly assessed risk-taking behaviors by incorporating ‘Time spent outdoors in winter/summer’ (Field IDs: 1060 and 1050), which reflects the potential risks linked to outdoor activities.

#### Substance use

We incorporated data on various smoking and drinking behaviors, alongside cessation experiences, to investigate the impact of neuroticism gradients on these habits. For smoking behaviors, we included ‘Ever smoked’ (Field ID: 20160, binary data) and ‘Current tobacco smoking’ (Field ID: 1239, categorized as ‘Yes, on most or all days’ = 1, ‘Only occasionally’ = 0.5, ‘No’ = 0). Given the adverse effects of smoking, promoting cessation among frequent smokers is a critical health strategy. However, the available data on cessation experiences, such as ‘Smoking compared to 10 years previous’ or ‘Ever stopped smoking for 6+ months’, exhibit significant bias. These data were collected exclusively from participants who reported past frequent smoking (26.4% of the sample). Additionally, this past smoking status was not collected from those who currently smoke daily (92.3% of daily smokers). To address this issue, we introduced a ‘quit heavy smoking’ metric, categorizing current heavy smokers as 1 (those who responded ‘Yes, on most or all days’ to ‘Current tobacco smoking’) and past heavy smokers as 0 (those who responded ‘Only occasionally’/’No’ to ‘Current tobacco smoking’ and ‘Yes, on most or all days’ to ‘Past tobacco smoking’). For drinking behaviors, we included variables ‘Alcohol intake frequency’ (Field ID: 1558), ‘Average weekly red wine intake’ (Field ID: 1568), and ‘Average monthly spirits intake’ (Field ID: 4440). The variable ‘Alcohol intake versus 10 years previously’ (Field ID: 1628) was utilized as a proxy for changes in drinking behaviors over time.

#### Physical activity

Existing literature consistently demonstrates that individuals with high levels of neuroticism are less likely to engage in physical activity^92,93^. We categorised the intensity of physical activity into three different levels: low, moderate and high, based on the ‘Summed MET minutes per week for all activity’ (Field ID: 22040). Specifically, total physical activity was classified as low (less than 600 MET minutes per week), moderate (600 to 3000 MET minutes per week), and high (more than 3000 MET minutes per week) in accordance with the established literature on the classification of total physical activity levels. In addition, our analysis included examination of sedentary behaviors commonly observed in adults. These included ‘Time spent watching television’ (Field ID: 1070) and ‘Weekly usage of mobile phone in last 3 months’ (Field ID: 1120).

#### Life satisfaction

Individuals with high levels of neuroticism typically exhibit lower levels of well-being^94,95^. We examined a range of aspects including ‘Happiness’ (Field ID: 4526), ‘Health satisfaction’ (Field ID: 4548), ‘Friendship satisfaction’ (Field ID: 4570), ‘Work/job satisfaction’ (Field ID: 4537), ‘Family relationship satisfaction’ (Field ID: 4559) and ‘Financial situation satisfaction’ (Field ID: 4581).

#### Sleep

To provide a nuanced assessment of sleep quality, we integrated four critical sleep factors: optimal sleep duration, insomnia, snoring and excessive daytime sleepiness, which were used together to calculate a comprehensive healthy sleep score. To assess optimal ‘Sleep duration’, individuals reporting 7-8 hours of sleep were assigned a score of 1, while all other durations were assigned a score of 0 (Field ID:1160). Insomnia was assessed by responses to ‘Sleeplessness/insomnia’ (Field ID: 1200), with ‘Sometimes’ and ‘Usually’ indicating the presence of insomnia and ‘Never/rarely’ indicating its absence. ‘Snoring’ was assessed directly from participants’ responses (Field ID: 1210), and for ‘Daytime dozing / sleeping’, individuals who reported dozing off or falling asleep during the daytime unintentionally and responded ‘All of the time’ or ‘Often’ to the relevant question (Field ID: 1220) were identified as having narcolepsy, while all others were classified as not having narcolepsy.

#### Reactions to suicide/stress

For suicidal thinking, we included indicators of suicidal behavior such as ‘Ever attempted suicide’ (Field ID: 20483), ‘Recent thoughts of suicide or self-harm’ (Field ID: 20513), and ‘Attempted suicide in past year’ (Field ID: 20484). For stress reactions, we assessed distress in the past month due to stressful events through three specific sensations: ‘Avoided activities or situations because of previous stressful experience in past month’ (field ID: 20495), ‘Repeated disturbing thoughts of stressful experience in past month’ (Field ID: 20497), and ‘Felt very upset when reminded of stressful experience in past month’ (Field ID: 20498).

We also investigated the associations between the neuroticism gradients and dietary habits, socio-economic status, and other variables of interest, including parental age at death, household income, qualification, and intelligence. The methodologies and outcomes have been documented in the supplementary materials (refer to Supplementary Table 14, Supplementary Notes and Supplementary Fig. 4). To conduct statistical analyses on the relationship between the neuroticism gradients and lifestyle factors, we used GLM through Python package StatsModels. Control variables of age, gender, and the interaction between age and gender were included in the analysis.

### Structural MRI Data Acquisition and Analysis

The high-resolution T1-weighted images were acquired using a Siemens Skyra 3T system (Siemens Healthcare, Erlangen, Germany), equipped with a standard 32-channel head coil. The acquisition process employed a magnetization-prepared rapid gradient-echo (MPRAGE) sequence, with the following parameters: repetition time set to 2000 ms; echo time at 2.01 ms; the capturing of 208 sagittal slices; a flip angle of 8°; a field of view (FOV) of 256 mm; a matrix size of 256×256; and a slice thickness of 1.0 mm, resulting in a voxel size of 1×1×1 mm. Detailed information regarding this imaging protocol is available at http://www.fmrib.ox.ac.uk/ukbiobank/protocol/V4_23092014.pdf.

To reveal voxel-wise regional gray matter volume linked to neuroticism phenotypes, our study adopted VBM^91,96^ analysis. This approach was implemented using the Computational Anatomy Toolbox 12 (CAT12^97^), devised by the structural brain mapping group at the University of Jena Hospital and incorporated into the SPM12 framework at the Institute of Neurology, London. Based on CAT12, we processed the T1-weighted images to correct for field inhomogeneity and segmented them into gray matter, white matter, and cerebrospinal fluid. This segmentation was further refined through an adaptive maximum a posteriori segmentation approach^98^ and enhanced by partial volume estimation^99^. Spatial normalization of these images was conducted using the ‘Diffeomorphic Anatomical Registration using Exponentiated Lie algebra’ (DARTEL) algorithm^100^. Modulated grey matter images were spatially smoothed with a 4 mm full width half maximum (FWHM) Gaussian kernel. After a thorough review of the preprocessed scans for any artifacts and ensuring the availability of neuroticism questionnaires at the 2.0-time point, a cohort of 30,221 individuals was remained for the VBM analysis. The VBM maps of these individuals were averaged, with a threshold of 0.1 established to define gray matter regions. Analyses were then focused on these specified areas.

In addition to voxel-based analysis, we also performed a ROI-based analysis for a better explanation (Supplementary Table 8). The human Brainnetome Atlas^101^ (https://atlas.brainnetome.org/bnatlas.html) was employed to provide extensive parcellation information, spanning a multitude of brain areas including the cerebral cortex, thalamus, basal ganglia, amygdala, and hippocampus, amounting to a total of 123 regions per hemisphere. The Human Brainnetome Atlas is a fine-grained, cross-validated atlas, constructed based on in vivo connectivity architecture. In complementing this, for cortical regions, we also applied the classical Brodmann parcellation^102^ based on cytoarchitecture information. Given the growing emphasis on the cerebellum in emotion processing and psychiatric disorders, we utilized a probabilistic atlas^103^ for cerebellar lobules and nuclei, delineating 17 subregions per hemisphere.

We employed a GLM model to elucidate the relationship between neuroticism gradients and regional gray matter volume. The model was formulated as *Y* = *β*_0_ + *β*_1_ × *X* + *c* × *Z* + *ε*, where Y denotes the regional gray matter volume (voxel or ROI level), X denotes the gradients of neuroticism, and Z represents a set of covariates, including sex, age, the interaction of age and sex, head size scaling, imaging center, and scanner table position. To concisely display the significant voxels in the voxel-wise analysis presented in Fig. 4a, we applied FDR correction at *P* < 0.001 for ERIS, acknowledging its stronger effect, and *P* < 0.05 for neuroticism, both with a cluster size threshold exceeding 100. For ROI-based analysis, the FDR-adjusted *P* values are provided. Considering the potential association between neuroticism and the second gradient with emotional circuits, we expanded our analysis to encompass the bilateral habenula (based on a recent thalamic atlas^104^; for neuroticism: right habenula, standardized *β* = -0.006, *P* = 0.26; left habenula, standardized *β* = -0.014, *P* = 0.015; for ERIS: right habenula, standardized *β* = 0.027, *P* = 9.9×10^-7^; left habenula, standardized *β* = 0.033, *P* = 6.1×10^-9^), basal forebrain (based on Julich-Brain cytoarchitectonic atlas^105^; for neuroticism: right Ch 4, standardized *β* = 0.001, *P* = 0.83; left Ch 4, standardized *β* = 0.004, *P* = 0.42; for ERIS: right Ch 4, standardized *β* = 0.018, *P* = 5.7×10^-5^; left Ch 4, standardized *β* = 0.026, *P* = 1.7×10^-8^), and BNST (based on Neudorfer et al^106^; for neuroticism: right BNST, standardized *β* = 0.006, *P* = 0.24; left BNST, standardized *β* = -0.003, *P* = 0.60; for ERIS: right BNST, standardized *β* = 0.025, *P* = 3.6×10^-6^; left BNST, standardized *β* = 0.020, *P* = 0.00019).

### Functional annotation analysis

To enhance the functional interpretation of our VBM findings, we employed Neurosynth (https://neurosynth.org/), a meta-analytical platform that facilitates the automatic synthesis of statistical maps from over 14,000 function magnetic resonance imaging studies. This process is achieved through the extraction of high-frequency keywords. Considering the extensive array of term maps provided by Neurosynth, we narrowed our focus to cognitive and behavioral terms, adhering to methodologies established in a prior study. The selected terms represent a confluence of those listed in Neurosynth and the Cognitive Atlas – an open-access ontology in cognitive science, featuring an extensive compilation of neurocognitive terms. Our selection included terms directly related to neuroticism, such as ‘anxiety’, ‘stress’, and ‘emotion regulation’, as well as a diverse range of others including ‘selective attention’, ‘facial expression’, and ‘navigation’. The complete list of these terms is presented in supplementary table 15. For each term, we downloaded the corresponding association map in Montreal Neurological Institute (MNI) 152 space, thresholded at *P* < 0.01, FDR corrected. These maps provide a probabilistic value at each voxel, indicating the likelihood of observing significant activation associated with a given search term, compared to studies not utilizing that term. We next proceeded to extract the top 10% of voxels with the largest effect sizes from both the ERIS and neuroticism statistical maps. The absolute values of these selected VBM voxels were then subjected to a weighted summation with the term-based statistical maps from Neurosynth. Due to the variance in intensity across terms, normalization of the weighted sums was achieved by dividing them by the total value of the respective term map. In the analysis of ERIS, only positively correlated voxels were incorporated; for neuroticism, we focused exclusively on negatively correlated voxels, a decision informed by the predominant trend of negative associations, leading to the exclusion of the caudate. This method of spatial correlation analysis was instrumental in identifying terms with substantial spatial concurrence with our VBM results. To visually articulate the results, we employed a word cloud to depict the foremost 15 terms associated with ERIS and neuroticism, respectively.

### Genetic Analysis

Genome-wide genotyping was conducted on participants of the UK Biobank using two purpose-designed arrays: the UK BiLEVE Axiom Array for approximately 50,000 individuals and the UK Biobank Axiom Array for approximately 450,000 participants. Direct genotyping was followed by imputation using the Haplotype Reference Consortium and UK10K reference panels, expanding the dataset to include approximately 96 million variants. Our GWAS analysis was employed based on this imputed genetic dataset from the UK Biobank July 2017 release. Quality control was then implemented using the PLINK GWAS software package^107^, version 2.0.0. Variants were excluded if the minor allele frequency was below 0.1%, the imputation quality score was less than 0.8, or there were deviations from Hardy-Weinberg equilibrium (*P* < 1.00×10^-7^), resulting in 10,056,631 SNPs being retained for further analysis (excluding the X chromosome). Participants with more than 10% missing genotypes, those who had undergone imaging procedures, and individuals of non-European ancestry (either self-reported or inferred genetically) were excluded from the study. Finally, we identified 267,311 nominally unrelated individuals using methods similar to those previously described. Association analyses of the ERIS and classic neuroticism score were adjusted for demographic and technical variables, including age, sex, genotyping batch, and array, as well as 40 principal components to account for population stratification.

Utilizing reference linkage disequilibrium (LD) scores for individuals of European ancestry from the 1000 Genomes Project, we employed LDSC^35^ software version 2.0.0 to estimate Single nucleotide polymorphism (SNP) based heritability, genomic inflation factor, and the LD score regression intercept. To identify independent genomic risk loci and significant SNPs, we used the functional mapping and annotation of GWAS (FUMA^40^, version 1.5.2). FUMA discerns independently significant SNPs (*P* < 5.00 × 10^−8^ and LD *r*^2^ < 0.60), among which those independent at LD *r*^2^ ⩾0.10 are classified as lead SNPs. Candidate SNPs exhibiting LD *r*^2^⩾0.60 with a lead SNP delineated the boundaries of a genomic locus. Loci separated by less than 250 kb were merged. For ERIS, we identified 13 lead SNPs and 12 genomic risk loci; for neuroticism, 50 lead SNPs and 41 genomic risk loci were identified. Our analysis did not aim to test the prediction power of PRS for the respective phenotypes but rather to examine their association with mortality rates. Consequently, we generated personalized PRS scores for individuals included in our GWAS analysis using the ‘score’ function from PLINK (version 1.07), based on our pre-calculated GWAS summary statistics and applying various common *P*-value thresholds: 0.1, 0.05, 0.01, 0.005, 0.001, 0.0001, 0.00001.

### HARs-associated genes

HARs are genomic loci that are conserved across species yet exhibit elevated divergence in humans compared to other species. In our study, we utilized a dataset of 2,737 HARs, as referenced from a prior study that integrated findings from multiple publications^37^. We then computed gene-based p-values for GWAS summary statistics related to ERIS and neuroticism using the FUMA, identifying 310 and 850 associated genes, respectively, after FDR correction. Subsequent overlap with the GWAS gene set yielded 1,610 genes associated with HARs included in our analysis. To assess whether genes implicated in ERIS and neuroticism through GWAS were significantly enriched within the HARs-associated gene set, we conducted a nonparametric permutation analysis. The MAGMA^39^, implemented within FUMA, was utilized to calculate gene *P*-values from the summary statistics. We applied FDR correction to select significant gene sets for ERIS and neuroticism (*P* < 0.05). Specifically, for the first 100, 200, and 300 genes as well as all significant gene sets for ERIS (310 genes) and neuroticism (850 genes), permutation tests were performed after excluding overlapping genes. Subsequently, we randomly selected corresponding numbers of genes from all GWAS genes, repeating this process 10,000 times to calculate the frequency of overlap with HARs-associated genes, thus constructing a null model. We then evaluated whether the observed overlap of MAGMA-derived genes with HARs-associated genes exceeded that predicted by the null model. The statistical significance was ascertained by comparing the actual overlap percentage against that from 10,000 random iterations, resulting in a nonparametric *P* value.

Additionally, we utilized positional mapping genes from the FUMA to repeat this process. Based on their physical positions in the genome, we identified 149 and 524 non-overlapping genes for ERIS and neuroticism, respectively.

